# Fine-tuned repression of Drp1 driven mitochondrial fission primes a ‘stem/progenitor-like state’ to accelerate neoplastic transformation

**DOI:** 10.1101/2021.03.05.434102

**Authors:** B Spurlock, D Parker, MK Basu, A Hjelmeland, G Sajina, S Liu, GP Siegal, A Gunter, A Moran, K Mitra

## Abstract

The opposing processes of mitochondrial fission and fusion are emerging as crucial regulators of stemness. Gene knockout of the master regulator of mitochondrial fission, Drp1, prevents neoplastic transformation. However, stem/progenitor cells maintaining repressed mitochondrial fission are primed for self-renewal and proliferation. Here, we demonstrate that only fine-tuned repression of Drp1 establishes a ‘stem/progenitor-like state’ towards supporting carcinogen driven neoplastic transformation of keratinocytes, while more complete Drp1 repression prevents it. Only fine-tuned Drp1 repression maintains small networks of fused mitochondria to sustain a unique gene-expression profile with elevated stem/progenitor cell functional markers (Krt15, Sox2 etc) and their regulators (Cyclin E). Cells with such a mitochondria-primed state are slow cycling, susceptible to transformation, and when enriched by mild carcinogen exposure sustains elevated self-renewal/proliferation to form less differentiated tumors. Therefore, our data for the first time highlights a ‘goldilocks’ level of Drp1 repression that supports stem/progenitor state dependent neoplastic transformation.

## INTRODUCTION

Stem cells exhibit plasticity as the state of stemness is modulated by various intrinsic and extrinsic factors (Magee, Piskounova et al., 2012)(Folmes, Dzeja et al., 2012). Involvement of mitochondria in regulation of the stem cell state is complex (Lisowski, Kannan et al., 2018)(Khacho & Slack, 2017, Zhang, Menzies et al., 2018). The master regulators of mitochondrial fission and fusion processes are emerging as crucial regulators of both embryonic and adult stem cells (Khacho & Slack, 2017, Lisowski et al., 2018)(Spurlock, Tullet et al., 2020). The Dynamin Related Protein 1 (Drp1) is the master regulator of mitochondrial fission that breaks larger mitochondria into smaller elements (Friedman & Nunnari, 2014, Kageyama, Zhang et al., 2011). The effects of Drp1 driven mitochondrial fission is opposed by fusion between mitochondria driven by the Mitofusins and Opa1(Chen & Chan, 2017)(Schrepfer & Scorrano, 2016). Therefore, Drp1 repression allows unopposed mitochondrial fusion to sustain a hyperfused mitochondrial state. Enhanced mitochondrial fusion sustains stemness of certain adult stem cells of neural, germ line and mammary lineages (Khacho, Clark et al., 2016, Senos Demarco, Uyemura et al., 2019, Wu, Chen et al., 2019). Particularly, repression of Drp1 supports stemness and repression of Mitofusin or Opa1 inactivates stemness in adult mouse neural lineage (Iwata, Casimir et al., 2020, Khacho & Slack, 2017). The other extreme state, i.e. unopposed mitochondrial fission, is critical for achieving pluripotency of stem cells during reprogramming (Prieto, Leon et al., 2016). However, sustained mitochondrial fission reduces pluripotency of stem cells (Zhong, Cui et al., 2019). Thus, it can be speculated that a balance of timely fission and fusion of mitochondria may be critical for maintaining stem cell states.

In various tumors, the bulk tumor cell populations are maintained by the adult neoplastic stem cells (also called tumor initiating cells) as they self-renew, proliferate and differentiate (Magee, Piskounova et al., 2012). Drp1 has been linked to tumor formation in various cancer types (Nagdas, Kashatus et al., 2019, Serasinghe, Wieder et al., 2015, Tanwar, Parker et al., 2016, Tsuyoshi, Orisaka et al., 2020, Xie, Wu et al., 2015). Drp1 activation sustains neoplastic stem cells at least in the astrocytic lineage (Xie et al., 2015), and genetic ablation of Drp1 prevents neoplastic transformation (Serasinghe et al., 2015). In contrast, an elevated mitochondrial fusion state, sustained by mitofusin, can drive immortalization of neoplastic stem cells to support tumorigenesis in a Drosophila model (Bonnay, Veloso et al., 2020). We found that select neoplastic ovarian epithelial stem cells (marked by Aldh) have lower levels of mitochondrial fission and can convert to a state of elevated mitochondrial fission when self-renewal and proliferation is activated. Importantly, the cells with minimum mitochondrial fission are found within a neoplastic stem cell sub-population that has >10 folds self-renewal and proliferation ability compared to the other neoplastic stem cell subpopulation (Spurlock et al., 2019)(Spurlock et al., 2021b). Thus, we proposed, that a repressed mitochondrial fission state may prime neoplastic stem cells towards maximizing their potential of self-renewal and proliferation. Similar priming was demonstrated in a normal hematopoietic stem cell subpopulation that maintains Drp1 repression (Liang, Arif et al., 2020). Here, we investigated the causative role of Drp1 repression in mitochondria driven priming of a stem/progenitor-like like state in skin cells, given skin stem cells serve as an excellent model for adult stem cells (Fuchs, 2016). We established a transformed skin keratinocyte model that sustains abundant self-renewing/proliferating cells with attenuated Drp1 activity. Using genetic perturbation strategies and single cell analyses of the parental and the transformed keratinocyte model we demonstrate that only fine-tuned repression of Drp1 establishes a ‘stem/progenitor-like state’ towards supporting neoplastic transformation, whereas more complete Drp1 repression prevents it. Our data for the first time highlights a ‘goldilocks’ level of Drp1 repression that supports stem/progenitor-like state dependent neoplastic transformation.

## RESULTS AND DISCUSSION

### Establishment of a mild carcinogen transformed keratinocyte model that maintains abundant self-renewing/proliferating cells and attenuated Drp1 activity

Chemical carcinogens like 2,3,7,8-tetrachlorodibenzodioxin (TCDD) is widely used for transforming skin tissue and the keratinocyte HaCaT cells that models key aspects of skin regeneration (Wincent, Bengtsson et al., 2012)(Hao, Lee et al., 2012)(Ray & Swanson, 2004)(Der Vartanian, Quetin et al., 2019)(Boelsma, Verhoeven et al., 1999, Jung, Jung et al., 2016, Schoop, Mirancea et al., 1999). The HaCaT cells carry non-functional p53 (St John, Sauter et al., 2000), which provides the appropriate cellular context given Drp1 inactivation increases cell proliferation in the absence of active p53 (Mitra, Rikhy et al., 2012, Mitra, Wunder et al., 2009). We used both mild and strong dose of TCDD to uncover any possibility of mitochondria based priming of stemness during TCDD driven neoplastic transformation. TCDD driven carcinogenesis is caused by activation and upregulation of the aryl hydrocarbon receptor (AHR) (Mulero-Navarro & Fernandez-Salguero, 2016) (Leclerc, Staats Pires et al., 2021). Exposure to milder (T-1nM) and stronger (T-10nM) doses of TCDD causes a comparable increase in cell proliferation of HaCaT cells in a standard transformation protocol (Fig. 1A). The stable transformed colonies of TCDD-1nM (TF-1) and TCDD- 10nM (TF-10) maintain comparable upregulation of AHR, even in the absence of TCDD (Fig. 1B). Unlike the parental HaCaT cells (Parental), both transformed cells are able to form subcutaneous xenograft tumors, confirming their transformation status (Supplementary. Fig. 1A). Interestingly, pathological evaluation of H&E stained tumor sections revealed that the TF-1 cells gave rise to malignant tumors harboring less differentiated (primitive) squamous cells with large nuclei (blue) and little visible cytoplasm (pink) (Fig. 1C, left). Whereas the TF-10 cells formed tumors harboring differentiated stratified squamous epithelium with cells having expansion of cytoplasm (Fig. 1C, right); the clear gaps represent artifacts secondary to cryo-sectioning.

**Figure 1.**
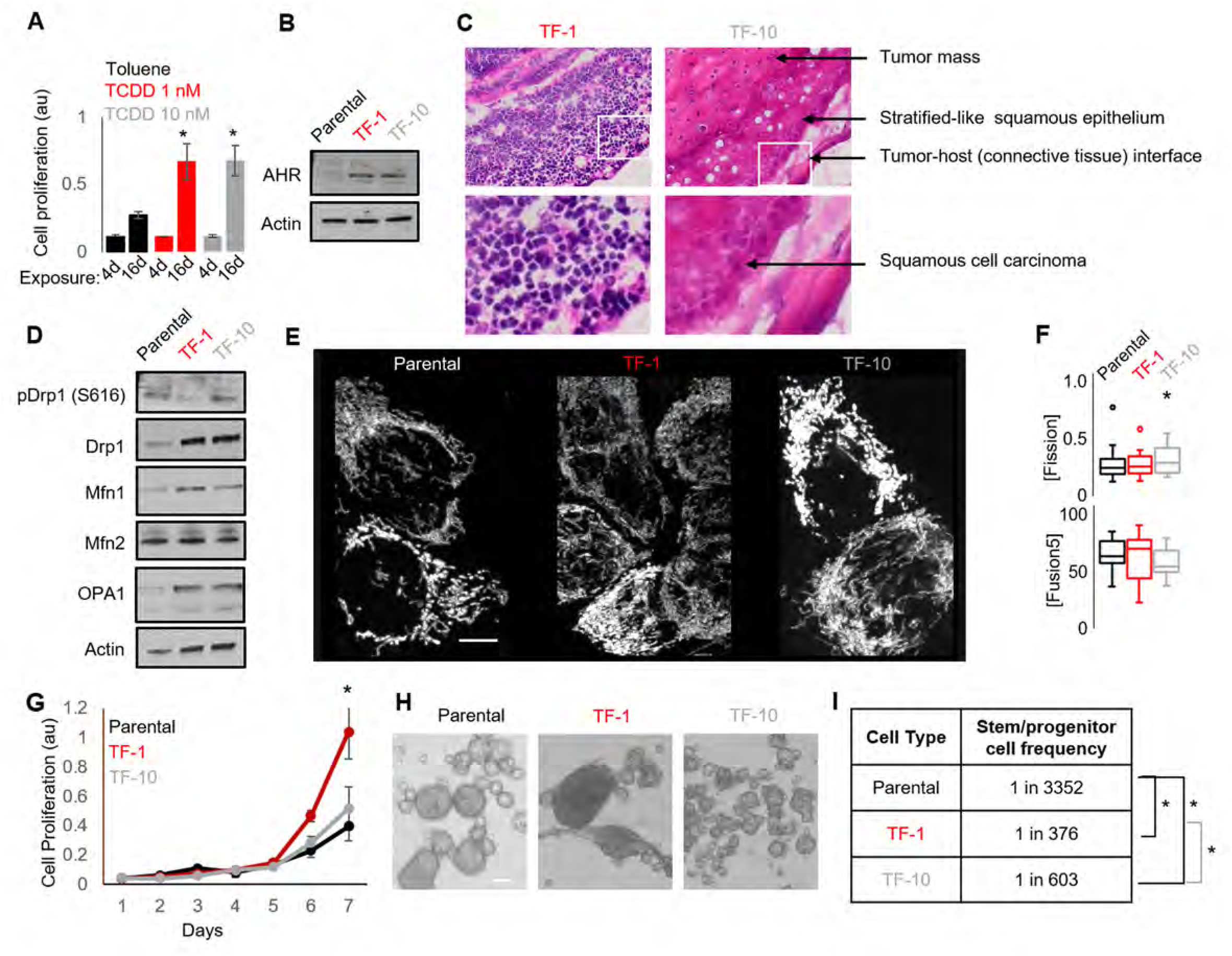
Establishment of a mild carcinogen transformed keratinocyte model that maintains abundant self-renewing/proliferating cells and attenuated Drp1 activity. **A)** Cell proliferation assay (quantified by crystal violet staining) of HaCaT cells after exposure to TCDD (1 or 10 nM) over 4 and 16 days, with Toluene (10nM) as vehicle control. **B)** Immunoblot analyses of AHR and actin (loading control) in transformed HaCaT populations, TF-1 and TF-10, in the absence of TCDD. **C)** Representative micrographs (40X) of H&E stained sections of tumor xenografts formed by TF-1 and TF-10 HaCaT cells; zoom of boxed region shown in the bottom panels. **D)** Immunoblot analyses of Drp1, pDrp1(S616), Mfn1, Mfn2, Opa1 and actin (loading control) in TF-1 and TF-10 HaCaT cells. **E)** Maximum intensity projection of confocal optical sections of representative Parental, TF-1 and TF-10 HaCaT cells expressing mitoPSmO. **F)** Quantification of mitochondrial [Fission] and [Fusion5] metrics from micrographs represented in (E). **G)** Quantification of cell proliferation assay (as in A) of Parental, TF- 1 and TF-10 HaCaT cells over 7 days in the absence of TCDD. **H)** Representative micrograph showing spheroids formed when 10^4^ Parental, TF-1 or TF-10 HaCaT cells were maintained in low attachment plate in presence of stem cell medium. **I)** Quantification of spheroid forming frequency of Parental, TF-1 or TF-10 HaCaT cells using ELDA. * signifies p value of <0.05 in T test (A,F) and ELDA (I); scale bar depicts 10 μm (D) and 100 μm (H).

Notably, among the major molecules regulating mitochondrial fission and fusion (Drp1, Mfn1, Mfn2, Opa1), Drp1 exhibited the most prominent differences (Fig. 1D). Particularly, the TF1 population has markedly lower levels of the cell cycle regulated activating phosphorylation of Drp1 (pDrp1-S616), while both the transformed populations have higher levels of Drp1 protein compared to the Parental line (Fig. 1D). The TF-1 population also has lower levels of other tested mitochondrial markers (Tom20, Cytochrome C and Fis1), compared to the TF-10 population (Supplementary Fig. 1B). Next, to assess the steady state contribution of mitochondrial fission and fusion on mitochondrial shape we quantified mitochondrial [Fission] and [Fusion5] metrics in individual live cells stably expressing the fluorescent mitochondrial reporter, mitoPSmO (Spurlock et al., 2019)(Spurlock et al., 2021b) (Fig. 1E). In comparison to the TF-1 and Parental populations, the TF-10 population exhibits reduced levels of the [Fusion5] metric and slightly elevated level of the [Fission] metric (Fig. 1F). Notably, although the TF-1 population has higher median [Fusion5], lower range of [Fusion5] is also detected in this population (Fig. 1F). Since the [Fusion5] metric reflects mitochondrial length in each cell (Spurlock et al., 2019), this data suggests a wider range of mitochondrial length in the TF-1 population, which could be potentially brought about by regulating Drp1 driven mitochondrial fission (Fig. 1D).

The TF-1 HaCaT population has significantly higher *in vitro* cell proliferation rate, compared to the Parental and the TF-10 population (Fig 1G). Furthermore, the TF-1 population form markedly larger spheroids in conditions that support self-renewal and proliferation, in comparison to those formed by the Parental or the TF-10 population (Fig. 1H). More importantly, ELDA statistics applied on spheroid formation assay for determination of *in vitro* stem cell frequency (Hu & Smyth, 2009) demonstrated that the TF-1 population has double the abundance of self-renewing/proliferating cells than the TF-10 population, while their abundance is one order higher in both transformed populations compared to the Parental (Fig. 1I).

The above data confirms the derivation of a transformed Drp1-attenuated-TF-1 keratinocyte model that maintains greater self-renewal/proliferative ability and forms less differentiated tumors compared to those transformed by a 10-fold higher TCDD dose. Thus, we conclude that the Drp1-attenuated-TF-1 keratinocyte population is enriched in neoplastic stem/progenitor cells.

### Transformed Drp1-attenuated-TF-1 keratinocyte population maintains an expanded slow-cycling sub-population with elevated stem/progenitor cell markers

To define the neoplastic stem cells of our newly derived transformed Drp1- attenuated-TF-1 population, we performed scRNA-seq (Luecken & Theis, 2019). Cell clustering in a UMAP plot shows 6 clusters within Parental, TF-1 and TF-10 populations (Fig. 2A, Clusters 0-5). While Cluster 4 is reduced in both transformed populations in comparison to the Parental, the TF-1 population exhibits a marked expansion of Cluster 3 and reduction of Cluster 5 (Fig. 2A, Supplementary. Fig.2A, left panel). These results also hold true with the lowest cluster resolution (Supplementary. Fig. 2B. arrows). We identified the top most candidate genes to mark each cluster (color coded arrows in Supplementary Fig. 2C, Supplementary. Data Table 1). Indeed, the neoplastic stem/progenitor cell enriched Drp1-att TF-1 population shows >3-fold upregulation of the epidermal stem cell marker Krt15 (Giroux, Lento et al., 2017, Gonzales & Fuchs, 2017, Purba, Haslam et al., 2014) and of the neoplastic stem cell markers for skin carcinomas, SOX2 and SOX4 (Boumahdi, Driessens et al., 2014, Foronda, Martinez et al., 2014) (Fig. 2B, arrow). Since the markers increase across all TF-1 clusters, with maximum levels in Cluster 3, we conclude that the TF-1 population exhibits upregulation of the stem/progenitor markers independent of their cluster distribution. On the other hand, the other stem/progenitor cell markers, like Krt5, Krt14, Krt19 (Gonzales & Fuchs, 2017)(Karantza, 2011) are also enriched by ∼2-fold in Cluster 3, but comparably in all the three cell populations, as exemplified by Krt5 (Fig. 2B, arrow, Supplementary Data Table 1). Notably, among the 3 cell populations, TF-1 harbors markedly fewer cells in Cluster 3 with elevation of the differentiating cell marker Krt13 (Karantza, 2011). Cluster 5, reduced in TF-1 population, is marked by 1.5 to >2- fold reduced expression of 12 of the 13mtDNA genes in all the 3 cell populations (Supplementary Fig. 2D, arrow). Cluster 4, reduced in both transformed populations, is marked by >8-fold upregulation of OASL, a gene involved in the interferon pathway, in all the three cell populations (Supplementary Fig. 2E).

**Figure. 2.**
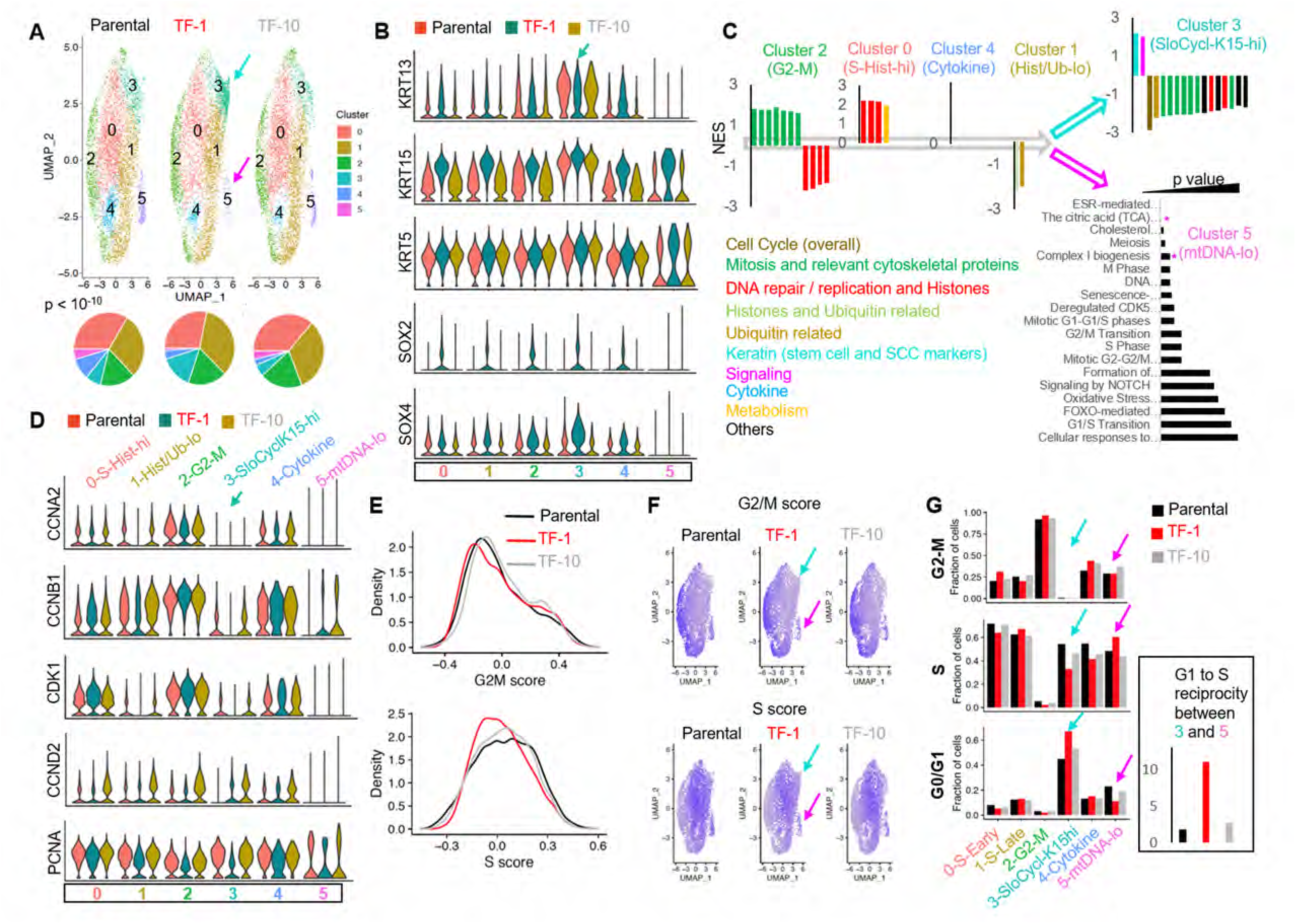
Transformed Drp1-attenuated-TF-1 keratinocyte population maintains an expanded slow-cycling sub-population with elevated stem/progenitor cell markers. **A)** UMAP plot of scRNA-seq derived clusters (0-5) of Parental, TF-1 and TF-10 HaCaT cells (top); pie chart of the percentage distribution of the clusters (bottom, p value from ChiSquare test); color coded arrows point the clusters with unique reciprocal abundance in TF-1 cells. **B)** Violin plots depicting expression of Krt13, Krt15, Krt5, Sox2 and Sox4 in the clusters identified in (A) in the Parental, TF-1 and TF-10 HaCaT cells; color coded arrow point to Cluster 3 marked by Krts. **C)** Normalized Enrichment Scores (NES) of functional pathways identified by GSEA analyses of marker genes of Clusters 0-4 (Supplementary data Table 2) and by overrepresentation analyses for Cluster 5 (* represents categories including mtDNA genes); functional categories defined by majority of the leading-edge genes are in parentheses, while clusters are arranged based on trajectory analyses (open arrows) (see Supplementary Fig. 2F). **D)** Violin plots depicting expression of cyclinA2, cyclinB1, CDK1, cyclinD2 and PCNA in the clusters identified in (A) in the Parental, TF-1 and TF-10 HaCaT cells; the functional category of genes identified in (C) is included in the cluster identity. **E)** Distribution of G2-M and S scores obtained by gene expression analyses of cell cycle predictive genes in the Parental, TF-1 and TF-10 HaCaT cells. **F)** Feature plot of G2-M and S scores obtained from (E). **G)** Bar plot showing fraction of cells in G2-M, S and G0/G1, computed from (E) in the functionally categorized clusters in the Parental, TF-1 and TF-10 HaCaT cells; inset shows G1 to S reciprocity of the clusters (color-coded arrows) with unique reciprocal abundance in TF-1 cells in (A).

Since flow cytometry on PI stained cells revealed modest differences in cell cycle distribution between the Parental and transformed populations (Supplementary. Fig. 2A, right panel), we asked if the scRNA-seq derived clusters represent distinct cell cycle phases. We performed the standard gene set enrichment analyses (GSEA) of the overall cluster marker profiles (Subramanian, Tamayo et al., 2005), where some statistically significant REACTOME pathways (q value <0.01) were renamed based on their leading-edge genes (Supplementary. Data Table 2). GSEA showed upregulation of genes in “Mitosis and Relevant Cytoskeletal Proteins” in Cluster 2 and that of genes in “DNA repair / replication and Histones” in Cluster 0 (Fig. 2C); Cluster 1 has downregulation of “Histones and Ubiquitin related” genes. Indeed, the mitotic Cyclin B and its partner kinase CDK1, as well as Cyclin A are markedly higher in Cluster 2-G2-M, while PCNA, which peaks early in S phase (Maga & Hubscher, 2003, Zerjatke, Gak et al., 2017) is highest in Cluster 0-S-Histone-hi in all 3 cell populations (Fig. 2D); the S phase cyclins E1/E2 and their partner CDK2 were not detected in our sc-RNAseq data set. Surprisingly, Krt15 marked Cluster 3 has dramatic downregulation of genes in “Overall Cell Cycle” as well as the above categories representing distinct cell cycle phases (Fig. 2C), suggesting their cell cycling is slow. Cluster 3-SloCycl-Krt15-hi maintains extreme low levels of major cyclins in all three cell populations, most prominently in TF-1 (Fig. 2D). PCNA, which is dramatically reduced in quiescent cells (Maga & Hubscher, 2003, Zerjatke, Gak et al., 2017), is markedly reduced in the SloCycl-Krt15-hi cluster (and others) only in the TF-1 population (Fig. 2D). SloCycl-Krt15-hi has upregulation of genes involved in “Keratin” and “Signaling” pathways (Fig. 2C). Notably, the Notch pathway target, Hes1, which actively maintains quiescence (Sang, Coller et al., 2008)(Moriyama, Durham et al., 2008), is upregulated across the clusters in the TF-1 population (Supplementary Fig. 2E). These data strongly suggest that the enrichment of the SloCycl-Krt15-hi cluster in the TF-1 population happens due to active maintenance of quiescence, rendering them slow cycling. Cluster 4 does not relate to cell cycle and shows upregulation of genes related to “Cytokine” (interferons) pathway (Fig. 2C), many of which are suppressed by AHR activation (Di Meglio, Duarte et al., 2014) (Supplementary. Data Table 1). This finding is consistent with our findings in Fig. 1B and Fig. 2A. The downregulated mtDNA genes in Cluster 5 were identified as “TCA cycle” and “Complex 1 biogenesis” in our overrepresentation analyses (* in Fig. 2C), but not in GSEA. Towards understanding the entry into the slow cycling state of the Krt15hi-Cluster 3, we performed trajectory analyses using Slingshot algorithm that identifies clusters related to each other based on their gene expression (Street *et al*, 2018). The output trajectory was the following: “G2-M” to “S-Histones-hi” to “Cytokine” to “Histones/Ub-lo” bifurcating into “SloCycl-Krt15-hi” or “mtDNA-lo” (Fig. 2C, open arrows, Supplementary. Fig. 2F). This implies that the cells in the TF-1 population may preferentially reside in the SloCycl-Krt15-hi cluster and not in the mtDNA-lo cluster, thus expanding the former and reducing the latter (Fig. 2A).

To confirm the cell cycle pathway results from GSEA, we quantified cell cycle scores and distribution in each cluster in cell population using a Seurat algorithm that employs several validated predictive cell cycle genes (Tirosh *et al*, 2016). This showed that the TF-1 population has lower S score and comparable G2-M score with the Parental and the TF-10 populations (Fig. 2E). Indeed, Cluster 2 is abundant in G2-M cells, Cluster 0 (and 1) in S and Krt15 marked Cluster 3 is in G0/G1 (expected for quiescent/slow-cycling cells) (Fig. 2F,G). This data taken with the lower levels of Histone transcripts in Cluster 1 than in Cluster 0 (* in Fig. Supplementary 2C) suggest that Cluster 0 may be in early S phase while Cluster 1 in late S phase. We noted that the expanded 3-SloCycl-Krt15-hi cluster in the TF-1 population has significantly more cells in G0/G1 and less cells in S than the other two populations (Fig. 2F,G, arrows), which is opposite to the 5-mtDNA-lo cluster that is reduced in the TF-1 population (Fig. 2F,G, arrows). This G1 to S reciprocity between Cluster 3 and 5 (quantified from the distributions) is >5-fold in the TF-1 population with respect to the Parental and TF-10 population (Fig. 2G, inset). This suggests that altered cell cycle regulation in the TF-1 population may allow them to preferentially reside in the 3-SloCycl-Krt15-hi cluster and not in the 5-mtDNA-lo cluster. Such a regulation may involve attenuation of CDK1 driven Drp1(S616) phosphorylation (Fig. 1D) due to marked reduction of CyclinB/CDK1 particularly in the TF-1 population in late S phase (Fig. 2D), where decision for expansion of the slow cycling Krt15-hi cluster is likely made (Fig. 2C, Suppl Fig. 2F).

Thus, our scRNA-seq characterization reveals that our newly derived transformed Drp1-attenuated-TF-1 keratinocyte population, enriched with self-renewing cells, maintain an expanded sub-population of slow cycling cells expressing high levels of stem/progenitor markers (lineage specific: Krt15; general: Sox2 and Sox4), likely due to altered cell cycle regulation. Our data is consistent with the Krt15+ epidermal stem cells, which generate various epithelial cell types or contribute to epidermal carcinoma, being primarily slow cycling (Giroux et al., 2017, Gonzales & Fuchs, 2017, Purba et al., 2014)(Morris, Liu et al., 2004). Moreover, the higher *in vitro* cell proliferation rate of the Drp1-attenuated-TF-1 keratinocytes (Fig. 1G) could result from the higher clonogenic capacity of the Krt15-hi cells (Morris et al., 2004)(Seykora & Cotsarelis, 2011).

### As opposed to more complete Drp1 repression, fine-tuned Drp1 repression sustains a [CyclinE^hi^-Sox2^hi^-Krt15^hi^] state and accelerates carcinogen driven keratinocyte transformation

Repression of Drp1 driven mitochondrial fission elevates stem/progenitor cell markers in mouse embryonic fibroblasts (MEFs), and in cells of neural and ovarian epithelial lineage (Iwata et al., 2020, Parker et al., 2015). In the epidermal lineage, we observed marked increase in in Sox2 protein levels (but no other embryonic stem cell markers) in the Drp1 attenuated TF-1 population in comparison to the Parental and TF- 10 keratinocyte population (Fig. 3A, Supplementary Fig. 3A), corroborating the scRNA-seq data (Fig. 2B). Krt15 protein levels are elevated in both transformed populations in comparison to the Parental (Fig. 3A), unlike the cluster specific differences in transcript levels (Fig. 2B). Immunostaining based co-expression analyses of Sox2 and Krt15 in individual cells revealed that only a subpopulation of Drp1-attenuated-TF-1 population has elevation of both Krt15 and Sox2 (Fig. 3B, Supplementary. Fig. 3B). Similar analyses showed that the highest levels of Sox2 in the TF-1 population is sustained in cells with an intermediate level of Drp1 protein levels (Fig. 3C) and lower pDrp1(S616) levels (indicative of lower Drp1 activity) (Fig. 3D). Therefore, we investigated whether repression of Drp1 in the Parental cells may cause the elevation of Sox2 and/or Krt15 observed in the TF-1 population derived from the Parental cells. Therefore, we knocked down Drp1 expression in Parental keratinocytes with two validated Drp1 shRNAs (Parker et al., 2015, Tanwar et al., 2016), one weaker (W) and the other 5 folds stronger (S) almost mimicking Drp1 genetic ablation (Fig. 3E). Surprisingly, while weaker Drp1 knockdown elevates Krt15 levels, the stronger Drp1 knockdown fails to do so (Fig. 3E). On the other hand, both weak and strong Drp1 knockdown caused marked elevation of Sox2 in a graded fashion (Fig. 3E), consistent with Drp1 ablated MEFs (Parker et al., 2015) and neural stem cells (Iwata et al., 2020). Thus, unlike the stronger Drp1 knockdown, the weaker Drp1 knockdown maintains detectable Drp1 protein, and uniquely sustains a Sox2-hi/Krt15-hi sub-population characteristic of the naturally Drp1 attenuated TF-1 population enriched in neoplastic stem cells.

**Figure 3.**
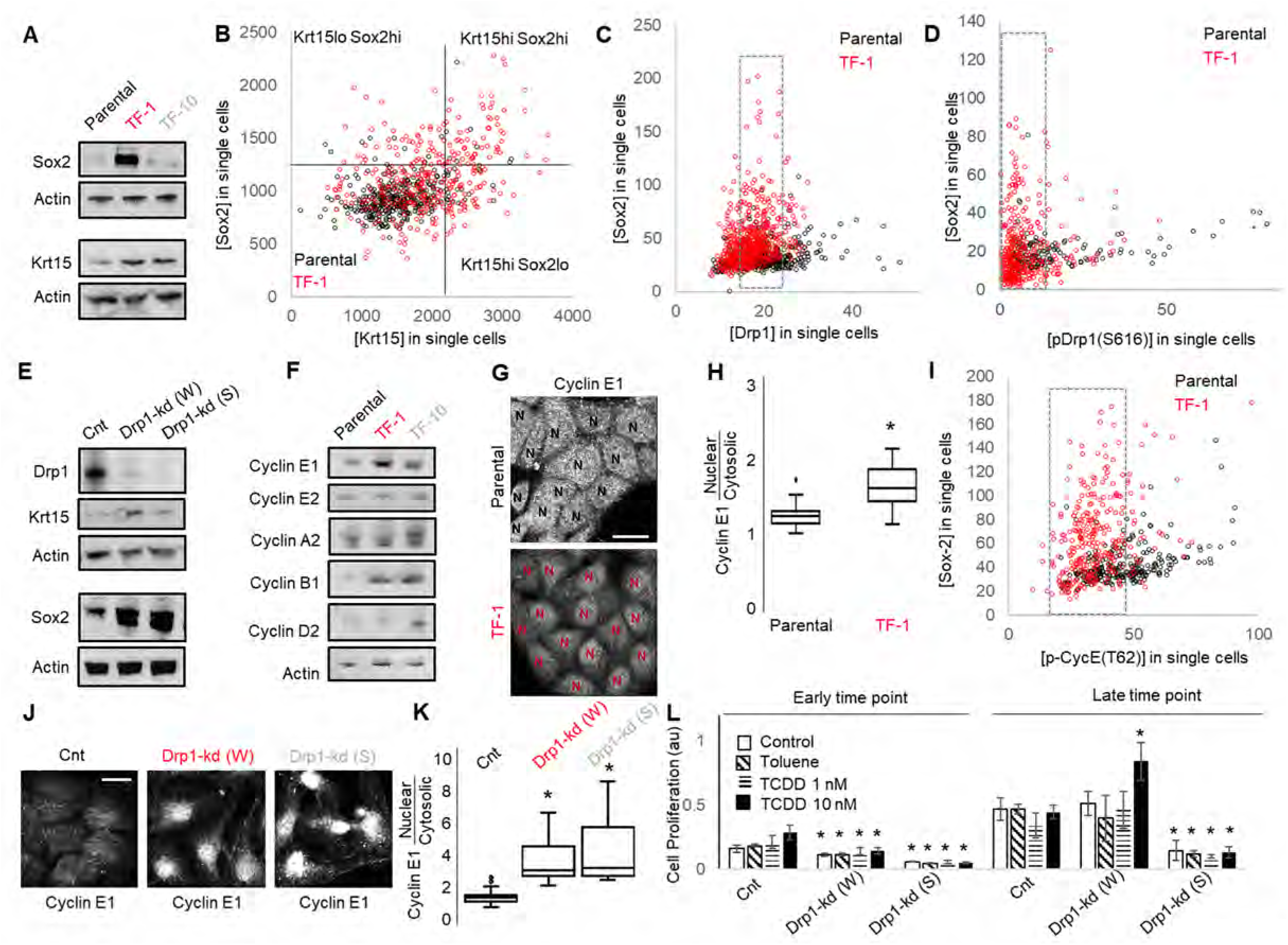
As opposed to more complete Drp1 repression, fine-tuned Drp1 repression sustains a [CyclinE^hi^-Sox2^hi^-Krt15^hi^] state and accelerates carcinogen driven keratinocyte transformation. **A)** Immunoblot analyses of Sox2, Krt15 and actin (loading control) in Parental and TF-1 and TF-10 HaCaT cells. **B)** Dot plot of [Sox2] and [Krt15] levels obtained from confocal micrographs of co-immunostained Parental and TF-1 HaCaT cells; black lines, marking 4 quadrants, represent levels of Sox2 and Krt15 determined from normalized signal in each population (see Supplementary Fig. 3B). **C)** Dot plot of [Sox2] and [Drp1] levels obtained from confocal micrographs of co-immunostained Parental and TF-1 HaCaT cells; dotted lines mark the levels of [Drp1] in cells expressing higher [Sox2] levels. **D)** Dot plot of [Sox2] and [pDrp1(S616)] levels obtained from confocal micrographs of co-immunostained Parental and TF-1 HaCaT cells; dotted lines mark the levels of [pDrp1(S616)] in cells expressing higher [Sox2] levels. **E)** Immunoblot analyses of Drp1, Krt15, Sox2 and actin (loading control) in Parental HaCaT cells expressing non-targeted shRNA (Cnt), weaker (W) or stronger (S) Drp1 shRNA. **F)** Immunoblot analyses of Cyclins E1, E2, A2, B1, D2 and actin (loading control) in Parental and stable TF-1 and TF-10 HaCaT cells. **G)** Representative confocal micrographs of Cyclin E1 immunostaining in Parental and TF-1 HaCaT cells; N depict the nucleus in each cell identified by DNA stain (not shown). **H)** Box plot showing quantification of nuclear to cytosolic ratio of Cyclin E1 levels quantified from experiment described in (G). **I)** Dot plot of [Sox2] and [CyclinE1(pT62)] levels obtained from confocal micrographs of co-immunostained Parental and TF-1 HaCaT cells; dotted lines mark the levels of [CyclinE1(pT62)] in cells expressing higher [Sox2] levels. **J)** Representative images of Cyclin E1 immunostaining in Parental HaCat cells expressing non-targeted shRNA (Cnt), weaker (W) or stronger (S) Drp1 shRNA. **K)** Box plot showing quantification of nuclear to cytosolic ratio of Cyclin E1 levels quantified from experiment described in (J). **L)** Cell proliferation assay of Parental HaCaT cells expressing non-targeted shRNA (Cnt), weaker (W) or stronger (S) Drp1 shRNA after exposure to TCDD (1nM or 10nM) at an early and a late time point in the neoplastic transformation protocol; ‘Toluene’ is the vehicle control for TCDD and ‘Control’ is with no chemical. * signifies p value of <0.05 in KW (G) and T test (L); scale bar depicts 20 μm.

Cell cycle status is a crucial determinant of stemness (Orford & Scadden, 2008, Otsuki & Brand, 2018, Pauklin & Vallier, 2013). Sox2 expression is under the influence of various G1 cyclins (Liu, Michowski et al., 2019). Cyclin E, which is indispensable for S phase entry from quiescence and also for neoplastic transformation (Geng, Yu et al., 2003, Siu, Rosner et al., 2012), is critical for sustaining Sox2 expression (Liu et al., 2019). Interestingly, among the major cyclins, Cyclin E1 levels are particularly higher in the Sox2 enriched Drp1 attenuated TF-1 population, while Cyclin D is particularly higher in the TF-10 population, when compared to the Parental, (corroborating scRNA-seq data, Fig. 2D) (Fig. 3F). Moreover, Cyclin E1 is preferentially accumulated in the nucleus in the TF-1 population, which is required to impact gene expression of Sox2 (and others) (Fig. 3G,H). The level of pCyclinE(T-62) reflects the level of active Cyclin E that drives transcription and remains susceptible to degradation (Siu et al., 2012). Immunostaining based co-expression analyses showed that in the TF-1 population, elevated Sox2 is sustained in cells with lower pCyclinE(T-62) than in the Parental population that has overall lower Sox2 (Fig. 3I). Moreover, cycloheximide chase assay, to assess degradation rate of proteins, revealed no Cyclin E degradation in TF-1 cells in comparison to marked Cyckin E degradation in the Parental cells; both populations maintain Tom20 (mitochondrial marker) within the 2 hour assay time frame (Supplementary. Fig. 3C). Thus, our data is consistent with elevated Sox2 in the TF-1 population being sustained by elevated Cyclin E levels achieved by attenuation of its degradation kinetics. Given mitochondrial regulation of Cyclin E regulates Cyclin E degradation (Mandal, Freije et al., 2010, Parker et al., 2015), we tested if repression of Drp1 driven mitochondrial fission in the Parental cells can elevate Cyclin E levels. Indeed, Cyclin E is dramatically accumulated in the nucleus in a graded manner with Drp1 knockdown efficacy (Fig. 3J,K), similar to Sox2 (Fig. 3E), suggesting Drp1 repression elevates nuclear Cyclin E and Sox2 levels similar to the naturally Drp1 attenuated TF-1 population.

Given, Sox2 and Cyclin E are important for neoplastic transformation in various cancer types (Hwang & Clurman, 2005, Schaefer & Lengerke, 2020, Teixeira & Reed, 2017), and Krt15+ cells give rise to skin carcinoma (Li, Park et al., 2013, Seykora & Cotsarelis, 2011), we asked if repression of Drp1 can modulate the process of carcinogen driven neoplastic transformation of keratinocytes. Thus, we exposed control and Drp1 knockdown Parental keratinocytes to 1 and 10nM TCDD following standard TCDD driven transformation protocol. Interestingly, the weaker Drp1 knockdown, which maintains Krt15-hi/Sox2-hi status and elevated nuclear Cyclin E, got transformed by 10nM TCDD even earlier than the control (Fig. 3L); 1 nM TCDD did not cause any transformation at this time point. This happened in spite of their lower proliferation rate than the control in the earlier time point. On the other hand, the stronger Drp1 knockdown, which also maintained slower cell proliferation rate and a Krt15-lo/Sox2-hi status with elevated nuclear Cyclin E, is not transformed in the same time window (Fig. 3L).

We conclude that the neoplastic stem/progenitor cell enriched Drp1-attenuated TF-1 population maintains a [CyclinE^hi^-Sox2^hi^-Krt15^hi^] sub-population. Fine-tuned Drp1 repression in the Parental cells primes the ‘stem/progenitor-like [CyclinE^hi^-Sox2^hi^- Krt15^hi^] state’, making them susceptible to neoplastic transformation, which is suppressed by more complete Drp1 repression. It is tempting to speculate that inability to sustain of this primed state may prevent neoplastic transformation in Drp1 ablated MEFs (Serasinghe et al., 2015).

### Fine-tuned Drp1 repression maintains smaller fused mitochondria, and sustains transcriptomic profile of the neoplastic stem cell enriched TF-1 population, which is suppressed by more complete Drp1 repression

Towards understanding how the ‘stem/progenitor-like state’ is primed by fine-tuned (incomplete) Drp1 repression, we performed detailed comparison of mitochondrial shape and over all gene expression between the Parental keratinocytes expressing control, weak or strong Drp1 shRNAs. Quantification of mitochondrial fission and fusion metrics contributing to mitochondrial shape in live cells confirmed that the stronger (more complete) Drp1 knockdown maintains ∼77% cells in the uniformly hyperfused state i.e. [Fission] metric <0.2 and [Fusion5] metric >80 (Spurlock et al., 2019), while the control population maintains ∼55% cells in this state (Fig. 4A, blue box, Supplementary. Fig. 3D). Interestingly, the weaker (incomplete) Drp1 knockdown maintains only ∼40% cells in the uniformly hyperfused state (Fig. 4A, blue box), while enriching a sub-population of cells (∼28%) in the [Fission] range of 0.2-0.4 and [Fusion5] range of 40-80 (Fig. 4A, red box), reflecting the mitochondrial fragments in the midst of extensive tubular mitochondria (Supplementary Fig. 3D). We also found that the cells in the enriched sub-population (Fission: 0.2-0.4 and Fusion5: 40-80) have lower mitochondrial number and total mitochondrial length (Fig. 4B), signifying decrease in mitochondrial content. However, cells with hyperfused mitochondria in the weaker Drp1 knockdown population (Fission<0.2; Fusion5>80) have distinctly higher mitochondrial number but not mitochondrial length (Fig. 4B). To confirm that such an increase in mitochondrial number is due to a greater number of smaller networks of fused mitochondria, we quantified the percent length of the longest mitochondrial elements (Fusion1). Indeed, less than 7% of cells have >80% of the mitochondrial length as hyperfused in the weaker Drp1 knockdown, compared to ∼40% in the stronger Drp1 knockdown (Fig. 4C, top). More importantly, the enriched population of cells in the weaker Drp1 knockdown with fission range of 0.2-0.4 have only up to 40% of the mitochondrial lengths in fused networks (Fig. 4C, bottom). We detected cells with similar mitochondrial fission and fusion features particularly in (FACS sorted) mitochondria primed ovarian neoplastic stem cell population that has 10-fold higher ability of self-renewal and proliferation (Spurlock et al., 2021b). We conclude that the remnant Drp1 driven mitochondrial fission after fine-tuned Drp1 repression enriches a subpopulation of cells maintaining smaller fused mitochondria (by preventing their hyperfusion) with reduced mitochondrial content and also distinct mitochondrial puncta, towards sustaining a [CyclinE^hi^-Sox2^hi^- Krt15^hi^] state. The functional significance of such previously unappreciated distinctions in mitochondrial shape may involve Drp1 driven regulation of mitophagy of damaged mitochondria (Twig & Shirihai, 2011)

**Figure 4.**
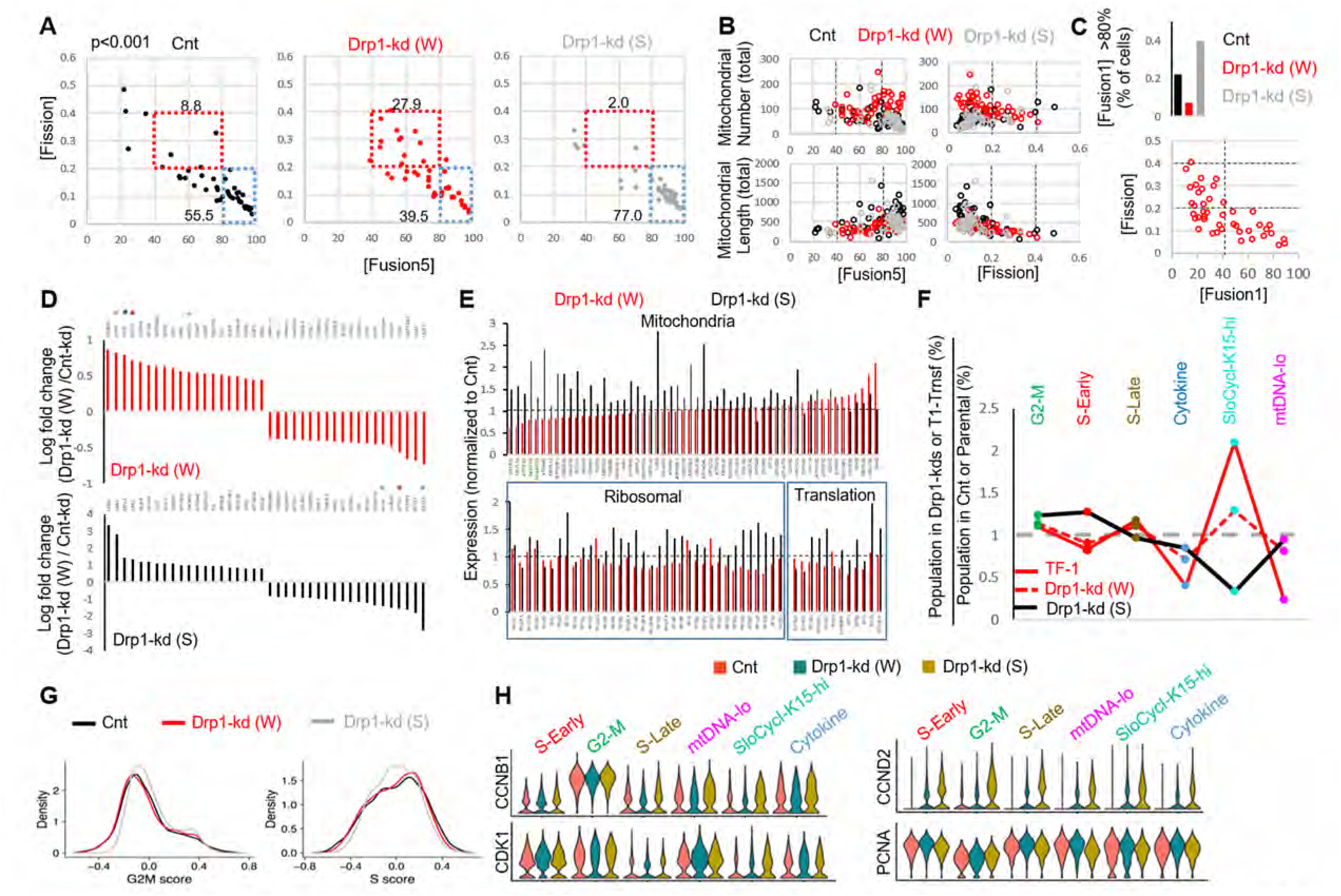
Fine-tuned Drp1 repression maintains smaller fused mitochondrial elements, and sustains transcriptomic profile similar to the neoplastic stem cell enriched TF-1 population, which is suppressed by more complete Drp1 repression. **A)** Dot plot of [Fission] and [Fusion5] obtained from confocal micrographs of Mitotracker green stained live Parental HaCaT cells expressing non-targeted shRNA as control (Cnt), weaker (W) or stronger (S) Drp1 shRNA; numbers denote the percentage of cells in the adjacent boxes with p value from ChiSquare test; representative images in Supplementary Fig. 3D **B)** Dot plot of [Fission] or [Fusion5] of single cells (from A) with their total mitochondrial number and total mitochondrial length; lines represent the ranges of [Fission] and [Fusion5] to demarcate cell subpopulations in boxes in (A). **C)** Bar graph signifying percentage of cells from (A) with >80 [Fusion1] metric (top); dot plot of [Fission] and [Fusion1] of cells from (A) (bottom). **D)** Barplots showing positive and negative log fold change values of top 20 genes in Parental HaCaT cells expressing weaker (W) (Top) or stronger (S) (bottom) shRNA with respect to the cells expressing non-targeting shRNA (Cnt); 0 signifies no change and color coded * denote relevant Krt genes. **E)** Barplot showing ratio of absolute expression values of mitochondrial genes (top) and ribosomal or protein translation genes (bottom) in the Parental HaCat cells expressing the stronger (S) or the weaker (W) Drp1 shRNA with respect to those expressing non-targeting shRNA (Cnt); 1 signifies no change. **F)** Line plot showing % population of Parental cells expressing the strong (S) or weak (W) Drp1 shRNA in the named scRNA-seq derived cell clusters, with respect to those expressing non-targeting shRNA (Cnt). This is compared to the same in the TF-1 cells with respect to its Parental population; 1 signifies no change from respective control. **G)** Distribution of G2-M and S scores obtained by gene expression analyses of cell cycle predictive genes in the Parental HaCaT cells expressing non-targeted shRNA (Cnt), weaker Drp1-shRNA (W) or stronger shRNA (S). **H)** Violin plots depicting expression of cyclinB1, CDK1 (left panel) and cyclinD2, PCNA (right panel) in the named scRNA-seq derived cell clusters in the Parental HaCaT cells expressing non-targeted shRNA (Cnt), weaker Drp1-shRNA (W) or stronger shRNA (S).

To probe the contribution of gene expression of any particular cell population in sustaining a stem/progenitor-like state after fine-tuned Drp1 repression, we performed scRNA-seq of the control and Drp1 knockdown Parental populations. Analyses of the overall gene expression (within the limits of gene coverage) confirmed that the weaker Drp1 knockdown, which maintains higher Drp1 transcript levels, modulates a smaller gene set to a smaller degree compared to the stronger Drp1 knockdown that maintains lower Drp1 transcript levels (Supplementary. Fig. 4A, B). However, graded Drp1 knockdown has both ‘similar’ (graded upregulation or downregulation) and ‘opposite’ effects (upregulation in one and downregulation in the other) (Supplementary Fig. 4C). The opposite effect is dominated by Krt15, Krt13, Krt4, Krt5 transcripts being upregulated in the weaker Drp1 knockdown and downregulated in the stronger Drp1 knockdown (color coded * in Fig. 4D, Supplementary Fig. 4C), corroborating our findings on Krt15 protein levels (Fig. 3E). On the other hand, genes for various mitochondrial proteins, involved in mitochondrial energetics, redox or biogenesis, were upregulated particularly in the stronger Drp1 knockdown (Fig. 4E upper panel, Supplementary Fig. 4C, bold). This is consistent with our observation of decreased mitochondrial content in the weaker Drp1 knockdown (Fig. 4B). Synthesis of the mitochondrial proteins can be potentially sustained by elevated ribosomal genes and genes involved in protein synthesis in the stronger Drp1 knockdown (Fig. 4E, lower panel). This finding is consistent with observations linking Drp1 repression to ribosomal genes and/or protein synthesis (Tanwar et al., 2016)(Zhao Q et al., 2021)(Favaro, Romanello et al., 2019). We speculate that repression of Drp1 beyond an optimal level may initiate compensatory changes through the retrograde pathway to trigger mitochondrial biogenesis (Ryan & Hoogenraad, 2007).

Cell clustering in a UMAP plot (Supplementary. Fig. 4D, Supplementary Data Table 3) revealed remarkable correspondence to the conserved cluster markers of the Parental/transformed data set (Fig.2, Supplementary. Fig. 4D, E), allowing assignment of cluster identities of G2-M, S-early, S-late, mtDNA-lo, SloCycl-Krt15-hi and Cytokine to the control and Drp1 knockdown data set. Strikingly, the weaker Drp1-knockdown population exhibits expansion of the SloCycl-Krt15-hi cluster and reduction of the Cytokine and mtDNA-lo clusters observed in the stem cell enriched Drp1-attenuated-TF-1 population, albeit to a lesser degree (Fig. 4F). This similarity in cluster organization suggests that fine-tuned Drp1 repression in Parental population establishes a stem/progenitor cell-like gene expression profile to poise them for accelerated neoplastic transformation (Fig. 3L). On the other hand, the stronger Drp1 knockdown markedly reduced the abundance of the SloCycl-Krt15-hi cluster and exhibits overall opposite trend of cluster profile compared to the weaker Drp1 knockdown and TF-1 population (Fig. 4F). Krt15 and various other marker Keratin genes are dramatically suppressed across the clusters in the stronger Drp1 knockdown and over all elevated in the weaker Drp1 knockdown (Supplementary Fig. 4F), confirming and expanding our findings on Krt15 protein levels (Fig. 3E). Moreover, analyses of cell cycle distribution (as in Fig. 2) revealed higher G2-M score, reduced S score and perturbed G1 to S reciprocity between SloCycl-Krt15-hi and the mtDNA-lo clusters particularly in the stronger Drp1 knockdown (Fig. 4G, Supplementary. Fig. 4G). The dramatic reduction of the SloCycl-Krt15-hi cluster in the stronger Drp1 knockdown cells could be due to their inability of exiting the elevated CyclinB1/CDK1 driven G2-M regulation in this cluster (Fig. 4H, left). Also, the stronger Drp1 knockdown maintained marked elevation of the growth factor cyclin, Cyclin D2 and reduction of the S phase marker, PCNA, across clusters (Fig. 4H, right); Cyclin E1 and Sox2 transcripts were not detected in this data set. Therefore, our data demonstrates that particularly fine-tuned Drp1 repression sustains a gene expression profile characteristic of the neoplastic stem cell enriched Drp1^-^attenuated-[CyclinE^hi^-Sox2^hi^-Krt15^hi^]-TF-1 population, which is prevented by more complete Drp1 repression that induces cell cycle disturbances.

Taken together, detailed molecular and mitochondrial shape analyses of our newly derived transformed TF-1 and its Parental keratinocyte model suggest, for the first time, that a fine-tuned “goldilocks” level of Drp1 repression maintains small networks of fused mitochondria to prime a slow cycling ‘stem/progenitor-like [CyclinE^hi^- Sox2^hi^-Krt15^hi^] state’ towards accelerating neoplastic transformation. Cyclin E, which maintains expression of stem cell markers, is indispensable for exiting cell cycle quiescence and also for neoplastic transformation (Hwang & Clurman, 2005, Schaefer & Lengerke, 2020, Teixeira & Reed, 2017)(Liu et al., 2019). We speculate that our defined CyclinE^hi^ state, enriched by fine-tuned Drp1 repression, may reduce depth of cell cycle quiescence (van Velthoven & Rando, 2019) towards rapidly transitioning to a self-renewing/proliferation state to support neoplastic transformation. Such transition may involve further activation of mitochondrial fission induced by signaling pathways supporting stemness which would be prevented with more complete Drp1 repression (as shown here) or total ablation of Drp1 (Kashatus, Nascimento et al., 2015, Serasinghe et al., 2015)(Spurlock et al., 2019). This conceptualization helps explain how regulated reduction and increase of Drp1 activity can potentially maintain adult stem cell properties (Iwata et al., 2020, Khacho & Slack, 2017, Parker et al., 2015, Xie et al., 2015). Further *in vivo* validation of our findings would reveal if and how bodily stresses causing mild repression of Drp1 driven mitochondrial fission, can possibly sustain the mitochondria primed ‘stem/progenitor-like state’ in tissues making them susceptible to neoplastic transformation. Enhanced mitochondrial fusion, proposed as a required event of neoplastic transformation of stem cells (Bonnay et al., 2020), may also involve such mitochondria driven priming.

## MATERIALS AND METHODS

### Materials

Biochemicals were obtained from Fisher Biochemicals or Sigma. Other materials: Drp1 shRNA (Dharmacon); Flouromount G (SouthernBiotech); FuGENE^®^ 6 Transfection Reagent (Promega); Luminata Forte Western HRP substrate (Millipore); 4% paraformaldehyde aqueous solution and Triton X-100 (FisherScientific); Labtek chambers (Nalgene Nunc International); DMEM (GIBCO); Geltrex (GIBCO); Primary antibodies were against: Drp1 (western blotting, 1:1000; immunofluorescence, 1:100; BD Biosciences); Cyclin E1 (HE12) (western blotting, 1:1000; immunofluorescence 1:100; Cell Signaling); actin AB-5 (1:10000; BD Biociences), Sox2 (western blotting, 1:500; immunofluorescence, 1:100; BD Bioscience), HSP-60 (1:200; BD Bioscience); Cytokeratin 15 (western blotting, 1:2000; immunofluorescence, 1:200; Abcam); Oct4 (immunofluorescence, 1:50; Cell Signaling); Nanog (immunofluorescence, 1:50; Cell Signaling). Secondary antibodies were from Jackson ImmunoResearch Laboratories. Athymic nude mice were purchased from Jackson laboratories.

### Cell culture methods

HaCaT cells were cultured on plastic dishes in Dulbecco’s Modified Eagle’s Medium (DMEM) with high glucose (4.5 g/L), sodium pyruvate (1 mM), L-Glutamine (4 mM), Penicillin (100 U/mL), Streptomycin (100 µg/mL), and 10% FBS using standard techniques. Transformed keratinocytes were derived from the Parental HaCaTs by selecting multiple clones after treatment with the noted dose of TCDD with replenishment with fresh TCDD every 2-3 days. The derived lines were treated with two courses of BM-Cyclin treatment to eliminate any mycoplasma infection.

Transfections were preformed using Fugene 6 Transfection Reagent modifying manufacturer’s protocols. Lines stably expressing Drp1-shRNAs or PSmO were selected using puromycin treatment (2 µg/mL) with regular media changes for 2 weeks until resistant colonies grew. The stable Drp1 knockdown lines were treated with two courses of BM-Cyclin treatment to eliminate any mycoplasma infection.

Standard cell proliferation assay was performed as described in (Parker et al., 2015). The extreme limiting dilution analysis (ELDA) of spheroid formation ability was performed by seeding 1, 10, 100, and 1000 cells per well with 24 wells per dose in a 96 well UltraLow Attachment plate and TIC-supporting media (DMEM:F12 supplemented with human growth factors as published elsewhere (Spurlock et al., 2019)). Wells were Supplementaryemented with media equaling 10% of the total volume every second day. On Day 5, the total number of wells for each dose containing nascent spheroids was tabulated. The online ELDA tool was used to analyze the results (Hu & Smyth, 2009).

### Immunoblotting and Immunofluorescence

Immunoblotting was performed using standard techniques. Whole cell lysates were run on 10% polyacrylamide gels and transferred to PVDF membranes followed by probing with appropriate primary and secondary antibodies.

Immunofluorecence was performed as described previously (Parker et al., 2015) on cells seeded in LabTek 8-well glass bottom or slide chambers. Briefly, cells were fixed in freshly prepared 4% aqueous paraformaldehyde supplemented with 4 %w/v sucrose and permeablized in freshly prepared 0.1 % Triton X-100 prior to staining.

Immunostained cells were mounted in Fluoromount with DNA stain Hoechst 33342 (10 µg/mL).

### Confocal microscopy, image processing and analysis

Confocal microscopy was performed on a Zeiss LSM700 microscope equipped with 405 nm, 488 nm, 555 nm, and 639 nm lasers, using the proprietary Zen Black (2012) software. Confocal micrographs were acquired with optimized laser powers and appropriate filters to minimize crosstalk, cross-excitation and bleaching. Live cells were imaged using a temperature and CO_2_ controlled chamber set to 37 C and 5% CO_2_.

Image processing and analysis of relative protein abundance and localization were performed using Zen Black and Zen Blue software to obtain background corrected mean fluorescence intensities within defined regions of interest drawn on maximum intensity z-projections of optical slices. Nuclear and cytosolic regions were demarcated based on DNA stain Hoechst 33342 and the immunostain of the molecule of interest, respectively. The [Fission] and [Fusion1/5] metrics using MitoGraph v2.1 software run on 3D stacks of confocal optical slices acquired from live cells expressing mito-PSmO or stained with Mitotracker Green, as described in (Spurlock et al., 2019). [Fission] is total mitochondria number / total mitochondrial length; [Fusion5] is (sum of top 5 mitochondrial length / total mitochondrial length) X 100. Microsoft Excel and IBM SPSS Statistics 23 were used to perform background corrections, bivariate analyses and significance testing.

### Tumor Forming Assay in mice and histochemistry

5 million transformed HaCaT cells were injected in subcutaneously in the flank of athymic nude mice to form xenograft tumors. Prior to injection, cells were trypsinized and washed once with sterile PBS, and finally resuspended to 25,000 cells/µL in room temperature PBS with Geltrex. Tumors arising within 6 to 8 months were harvested for histochemical analyses. The harvested tumors were frozen unfixed in TissueTek OCT compound on dry-ice. A Cryostat was used to obtain 5 µm slices, which were then fixed using freshly made cold 4% paraformaldehyde and then stained with H&E using standard techniques.

### Single Cell RNAseq and data analyses

Trypsinized single cells were washed in PBS (without ca++ or Mg++) with 0.04% BSA and tested for >90% viability. Single cell analysis was performed on a 10xGenomics platform according to 3’ v3.1 NextGem Dual Index manual. The 3’-biased cDNA libraries were constructed through the following steps: cDNA fragmentation, end repair & A-tailing, and size selection by SPRIselect beads, adaptor ligation, and sample index PCR amplification, and then SPRIselect beads size selection again. The final constructed 3’- biased single cell libraries were sequenced by Illumina Nextseq500 machine, targeting total reads per cell for 20,000 at minimum, and the sequencing cycles consisted of 28bp for read 1, 90 bp for read 2, and 10 bp for i7 index, and 10bp for i5 index.

Count matrices were generated from the single-cell raw fastq files using 10x genomics *cellranger* software (v.4.0) using hg38 reference genome provided by 10x Genomics. We filtered the data to only include cells expressing at least 2000 detectable genes (using “nFeature_RNA” filter); this also filtered out dead cells with >10% mt-DNA gene expression in our data set. Our data coverage included approximately ∼ 3000 genes in average of 5000 cells in each population. The resulting count matrices were analyzed by Seurat package (v3.9.9) with the standard workflow (Butler *et al*, 2018; Stuart *et al*, 2019). After applying the (*nFeature_RNA* > 2000), the expression data were then normalized using the *NormalizeData* function in Seurat and variable featuresets were identified using the *FindVariableFeatures* function in Seurat. Depending on the type of comparison, specific sets of samples were combined using the *IntegrateData* function. The integrated datasets were scaled and cells were clustered with up to 20 dimensions (dim=1:20). We performed clustering using several resolution parameters (0.1-0.8) and visually selected optimal resolution for specific datasets. The clusters were visualized using UMAP with up to 20 dimensions. Marker genes for each cluster were calculated with the *FindAllMarkers* function and statistical significance was calculated by Wilcoxon Rank Sum test. Differential expression of the cluster markers was carried out using the FindMarkers function of Seurat. For this, we only used the normalized data before integration (assay ‘RNA’) using the Wilcoxon rank sum test, as suggested by Seurat documentation. For cell-cycle scoring, we use the CellCycleScoring function of Seurat, which provides scores of G2M and S phase scores and assigns cell-cycle phases based on G2-M and S scores, or in G1/G0 when both scores are low (Tirosh *et al*, 2016). The cell-lineage determination and trajectory calculation were carried out using the *Slingshot* (Street *et al*, 2018) package with Seurat clusters and default parameters. The algorithm was allowed to automatically identify the start and the end clusters. GSEA analyses were performed with ranked log fold change (LFC) of markers (>0.1, p<0.01) using MsigDB as described in (Tanwar et al., 2016). Overrepresentation analyses were performed with genes with negative LFC in Cluster 5 using the Reactome PA package of R v 3.6.0.

### Statistical analyses

Drawing of plots and statistical analyses were performed using Excel, SPSS or R package as mentioned in appropriate section above.

## ACKNOWLEDGEMENTS

BS, KM, AM are supported by NIH award (# R33ES025662). We acknowledge Dezhi Wang from the Pathology Core Research Laboratory for preparation of H&E stained slides, UAB Flow Cytometry and Single Cell Services Core (supported by AR048311, AI027667) and the Dr. Mike Crowley from the Heflin Genomics Core for sc-RNAseq.

## AUTHOR CONTRIBUTION

BS designed and performed experiments and analyzed data along with AM and AG; DP generated the transformed lines and performed basic characterization; ABH supervised the xenograft experiments performed by BS and SG; GPS performed the pathological evaluation of tumor sections; SL provided consultation for the sc-RNAseq and prepared the scRNAseq library; MKB provided consultation for the sc-RNAsq analyses and performed the analyses; KM conceived the study, helped in experimental design, analyzed data and wrote the manuscript with BS.

## SUPPLEMENTARY FIGURE LEGENDS

**Supplementary Figure 1.**
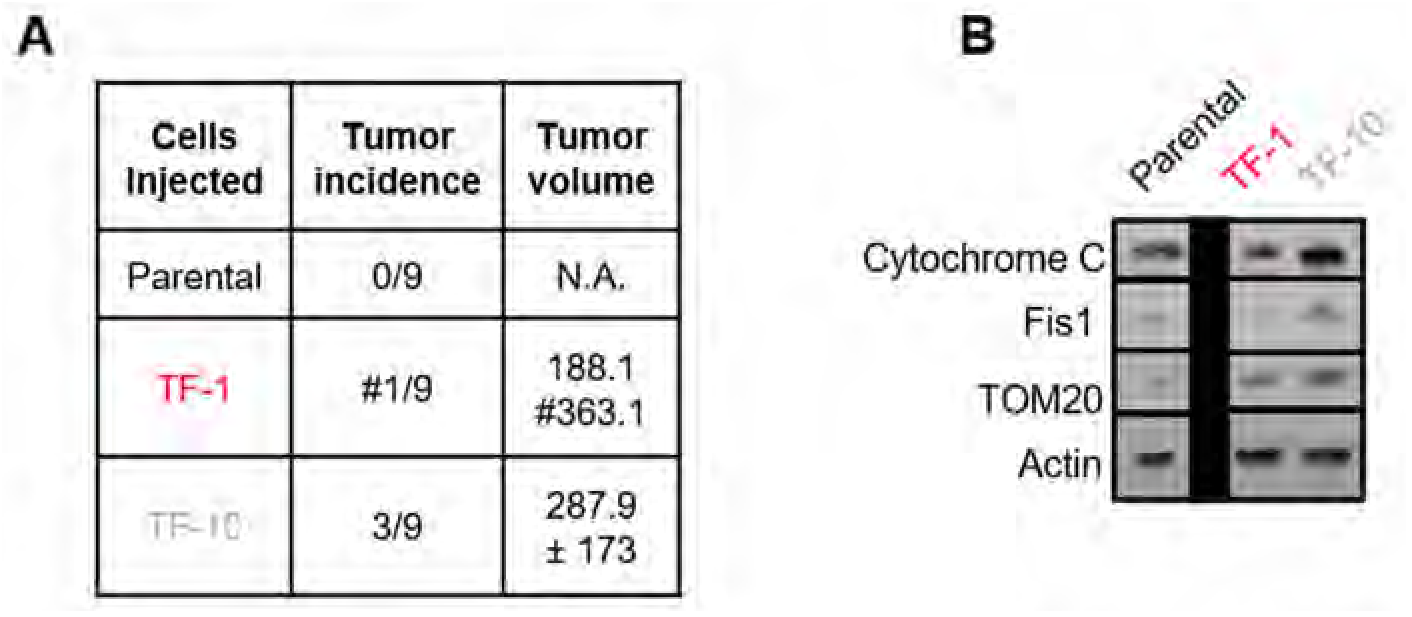
**A)** Quantification of incidence and volume of xenograft tumors formed by TF-1 and TF-10 HaCaT cells; # indicates presence of a distant tumor. **B)** Immunoblot analyses of Cytochrome C, Fis1, Tom20 and actin (loading control) in Parental and TF-1 and TF-10 HaCaT cells.

**Supplementary Figure 2.**
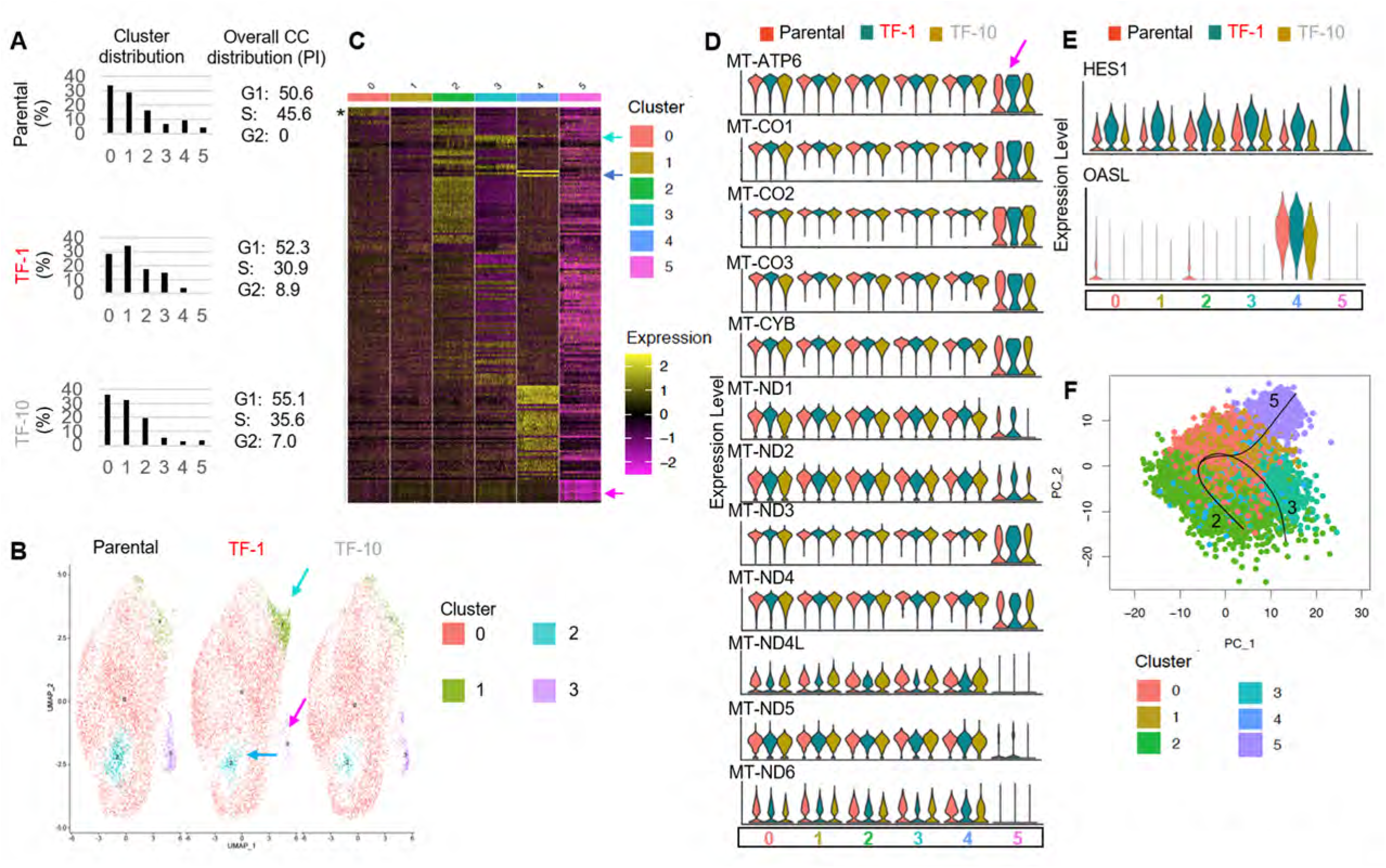
**A)** Bar plot showing the percentage distribution of the clusters (0-5) in each of the cell populations identified by scRNA-seq analyses of Parental, TF-1 and TF-10 HaCat cells (left); numbers show cell cycle distribution from flowcytometry analyses of PI stained cells in the corresponding population (right). **B)** UMAP plot of scRNA-seq derived clusters (0-5) of Parental, TF-1 and TF-10 HaCaT cells obtained with minimum cluster resolution; color coded arrows point the major change in cluster distribution in transformed cells. **C)** Heat map of statistically significant marker genes (LFC ≥1.5 folds, p-adjusted <0.0001) for the color-coded clusters (0-5) of Parental, TF-1 and TF-10 HaCaT cells; color coded arrows point to the top most marker gene in the 3 clusters that are have different distribution between the Parental and transformed populations; color index shows log fold change; * marks the Histone gene cluster. **D)** Violin plots depicting expression of the mt-DNA genes in the named scRNA-seq derived cell clusters in the Parental, TF-1 and TF-10 HaCaT cells. **E)** Violin plots depicting expression of HES1 and OASL genes in the named scRNA-seq derived cell clusters in the Parental, TF-1 and TF-10 HaCaT cells. **F)** PCA plot of trajectory analyses of the scRNA-seq derived cell clusters in the Parental, TF-1 and TF-10 HaCaT cells.

**Supplementary Figure 3.**
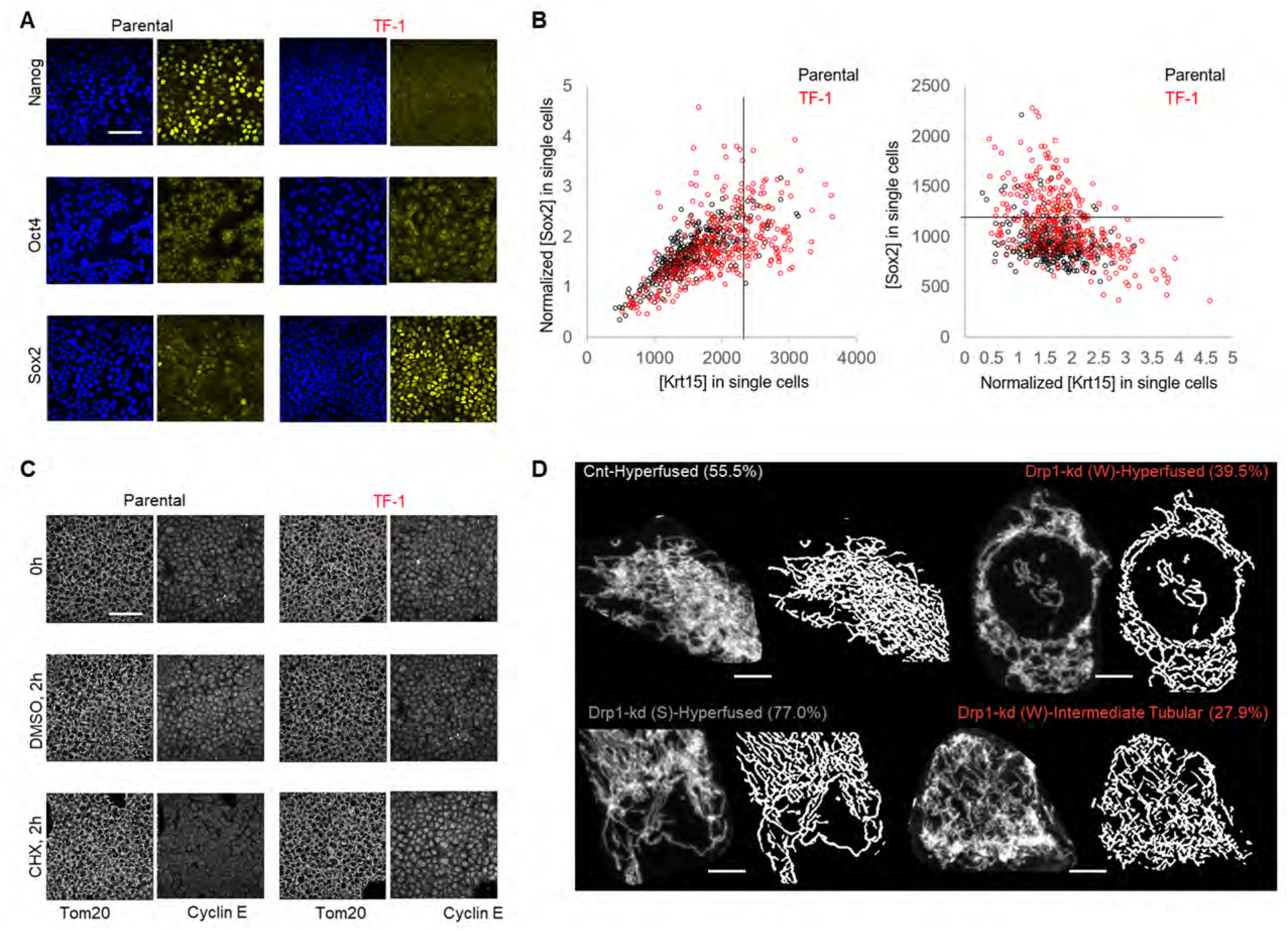
**A)** Representative confocal micrographs of immunostained Parental and TF-1 HaCaT cells showing endogenous expression of Oct4, Nanog and Sox2; blue depicting nuclear staining with Hoechst. **B)** Dot plot of [Sox2] and [Krt15] obtained from confocal micrographs of co-immunostained Parental and TF-1 HaCaT cells; [Sox2] levels normalized to the minimum in the respective population (left) and [Krt15] levels normalized to the minimum in the respective population (right); black lines demarcate levels of Sox2 and Krt15 as determined from the normalized signal of the other. **C)** Representative confocal micrographs of Parental and TF-1 HaCaT cells showing endogenous expression of Tom20 and Cyclin E1 after 2 hours incubation with DMSO or Cychloheximide, with the pre-incubation time point (0h) as control. **D)** Representative maximum intensity projections (MIP) of confocal optical sections (micrograph) and the corresponding MitoGraph generated binary images (binary) for quantifying [Fission] and [Fusion1/5] metrics of the Parental HaCaT cells expressing non-targeted shRNA (Cnt), strong (S) or weak (W) Drp1 shRNA; the cells represent populations in colored boxes in Fig. 4A while the number depicts their abundance in percentage. Scale bar depicts 100 μm (A, C) or 15 μm (D).

**Supplementary Figure 4.**
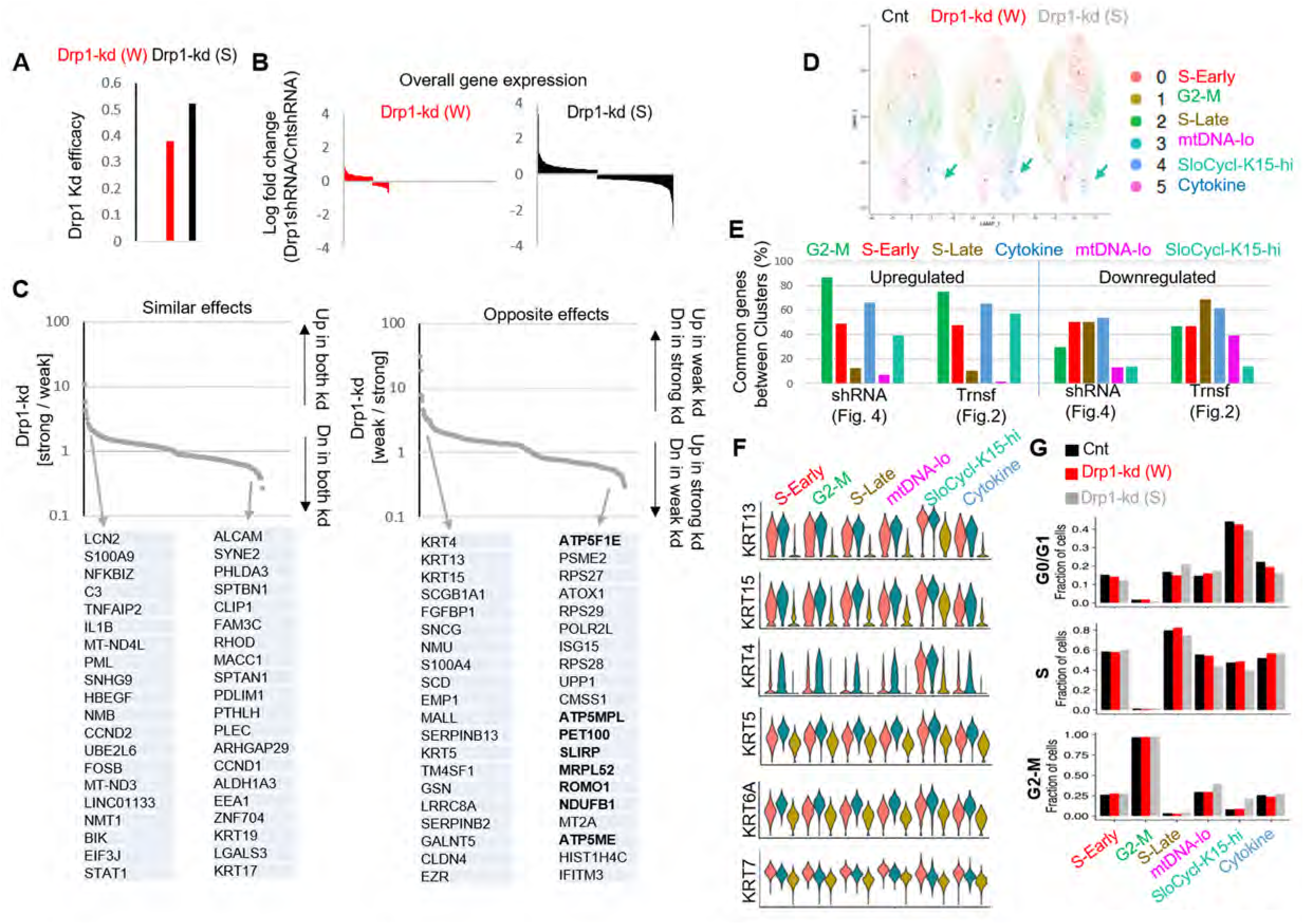
**A)** Bar plot showing the knockdown efficiency of strong (S) or weak (W) Drp1 shRNA expressed as ratio of Drp1 transcript levels in the knockdown HaCaT and that of the cells expressing the non-targeted shRNA control. **B)** Bar plot showing log fold change of overall gene expression of Parental HaCaT cells expressing strong (S) or weak (W) Drp1 shRNA with respect to those expressing the non-targeted shRNA (Cnt). **C)** Plots showing ratio of the expression of covered genes between the Parental HaCaT cells expressing strong (S) or weak (W) Drp1 shRNA; “Similar effects” showing ratio of increase or decrease in the (S) Drp1 shRNA over the (W) Drp1 shRNA (back arrows in left panel), while opposite effects showing opposite trends in genes as labelled in the arrows (back arrows right panel); grey arrows point to the top 20 genes of each category. **D)** UMAP plot of scRNA-seq based clusters (0-5) of Parental HaCaT cells expressing non-targeted shRNA (Cnt), strong (S) or weak (W) Drp1 shRNA; arrow pointing the SloCyc-Krt15-hi cluster (note color coding is not same as the one presented in Fig. 2). **E)** Bar plot showing percentage of statistically significant common genes (Upregulated or Downregulated) between color coded clusters obtained with the scRNA-seq data set of parental and transformed HaCaT cells (Fig. 2) and that of parental expressing control or Drp1 shRNAs (Fig. 4); same color coding is maintained between the two data sets. **F)** Violin plots depicting expression of Krt13, Krt15, Krt4, Krt5, Krt6A, Krt7 genes in the named scRNA-seq based cell clusters in the Parental HaCaT cells expressing non-targeted shRNA (Cnt), strong (S) or weak (W) Drp1 shRNA. **G)** Bar plot showing fraction of cells in G2-M, S and G0/G1, computed from main Fig. 4G, in the functionally categorized clusters in the Parental cells expressing non-targeted shRNA (Cnt), strong (S) or weak (W) Drp1 shRNA.

